# Biodiversity data portals as methodological filters for iNaturalist: how platform choice can shape biodiversity assessments

**DOI:** 10.64898/2026.06.12.731923

**Authors:** Elizabeth Edson, Diego Ellis-Soto, Avery P. Hill, Rebecca F. Johnson

## Abstract

iNaturalist has rapidly grown to become one of the largest contributors of global biodiversity data, being widely used in academic research, to support and inform applied conservation, and to help develop policy indicators for decision makers. However, the availability of iNaturalist data varies based on the digital platform it is accessed from. Here, we assess whether the pathway used to access iNaturalist data from three commonly available biodiversity data sources: iNaturalist, the Global Biodiversity Information Facility (GBIF) and ESRI’s ArcGIS Online, alters occurrence record availability and downstream analyses for ecological inference. First, we investigate iNaturalist data availability on ArcGIS Online and find that the reduced field metadata for record location and obscuration information can lead to biased assumptions in the spatial ranges of sensitive species. Second, when assessing iNaturalist records available through GBIF, we find that restrictive Creative Commons observation licenses prevent an average of 26.1% of iNaturalist Research Grade records from being exported to GBIF, and this can lead to differences in environmental niche analysis when compared to datasets including all available research grade records. Understanding differences in platform licensing when integrating across biodiversity data repositories is another consideration for researchers and practitioners when conducting biodiversity assessments. Our results show that the pathway through which iNaturalist data are accessed can function as a methodological filter, potentially altering spatial coverage, the climatic conditions represented by occurrence datasets, and downstream ecological analysis.

## 1. Introduction

Biodiversity, the variety of all living things on earth, is declining faster than at any time in human history (IPBES (2019)). In response to this ongoing biodiversity crisis, the 2023 Kunming–Montreal Convention on Biological Diversity adopted a Global Biodiversity Framework, stating its long-term vision to “live in harmony” with biodiversity by 2050 (CBD (2022)). This framework highlights the need for improved biodiversity monitoring and data collection, and efficient sharing of biodiversity information to support decision-making (Secretariat of the Convention on Biological Diversity (2024b)). Twenty-three targets for 2030 were identified in the Global Biodiversity Framework, including two that directly relate to biodiversity data: Target 14 aims to integrate biodiversity data in decision making at every level, and Target 21 aims to ensure that knowledge is available and accessible to guide biodiversity action (Secretariat of the Convention on Biological Diversity (2024a)).

A major challenge to Target 21 is aggregating and standardizing biodiversity information from different sources and formats. This has given rise to many different initiatives to build the data repositories and infrastructures necessary to support data-driven biodiversity research locally and globally. For instance, there are international initiatives such as the Global Biodiversity Information Facility (GBIF, www.gbif.org), and national initiatives such as the Atlas of Living Australia (https://www.ala.org.au/) and the UK’s National Biodiversity Network (https://nbn.org.uk/). These platforms are curators of primary biodiversity data, making standardized datasets freely available from a wide range of data providers such as natural history museum collections, individuals, public research institutions, and increasingly, participatory (citizen and community) science (Güntsch et al. (2024)). Other institutions such as ESRI host biodiversity datasets within broader content repositories like their Living Atlas (https://livingatlas.arcgis.com/en/home/), though biodiversity remains a minor, if growing, component of their wider geospatial portfolio of data.

Participatory science platforms, such as iNaturalist and eBird are fast becoming major contributors to both national and international biodiversity data repositories, because millions of participants are collecting large quantities of data at larger spatial and temporal scales, and at a lower cost, than traditional data collection methods. (Backstrom et al. (2024), Mason et al. (2025)). Robust data portals from international platforms such as GBIF and ArcGIS Online, have led to increased development of different pipelines of biodiversity data for integration and modeling on global and local scales. This easier integration of disparate sources of biodiversity information into decision making processes directly supports Target 14 of the GBF.

Since the launch of iNaturalist over a decade ago, it has rapidly grown to become one of the most widely used global platforms for biodiversity participatory science. iNaturalist is now a large repository of global biodiversity data and as of October 2025 had over 270 million observations from nearly 4 million observers (current numbers: https://www.inaturalist.org/observations). Between 2020 and 2023, about half of all the species in GBIF had over half of their records provided by iNaturalist (Loarie (2023)), highlighting the important contribution iNaturalist users have in biodiversity data collection. A large community of identifiers manages data quality through a process of identification consensus. Observations can only become “Research Grade” if they are a wild or naturalized organism, have a date, location coordinates, evidence (photo or sound), and two or more community members agree on the identification (iNatHelp (2025a)). Though some records will never make it to Research Grade, Cambell *et al*. showed in their paper “Identifying the Identifiers”, that nearly 60% of all iNaturalist observations do become elevated to Research Grade within a year of their recorded date (Campbell et al. (2023)), and thus become relevant for downstream data applications.

A combination of data quality assessment, ease of acquisition, large data volume, and global geographic coverage makes iNaturalist data extremely attractive for biodiversity research and decision making. The data has been utilized for research all over the globe with its use in peer-reviewed research growing 10-fold within the last 10 years; 1410 articles were published in 2022 alone that utilized iNaturalist data, with the most common focus being species distribution, niche modeling, and range dynamics (Mason et al. (2025)). iNaturalist data is also increasingly being used in non-academic research, such as during environmental reviews and environmental impact statements (Callaghan et al. (2025)).

Countries and sub-national governments that are working towards their Global Biodiversity Framework’s 30 x 30 targets (Secretariat of the Convention on Biological Diversity (2024a)), but do not have the support of a national database, may be more reliant on global databases and therefore participatory science data, to fulfil their local reporting needs. iNaturalist Networks (iNaturalist (2025)) allow some signatory countries to have a localized and branded iNaturalist portal which offers a better resolution of data within the network than is available to the general public, by providing the true coordinates of species observations to the government and approved non-profit partners such as local conservation institutions. A simpler, non-branded version of an iNaturalist Network is being developed for California, and this “California Data” will directly enhance State data repositories (California Academy of Sciences (2026)). In addition to being accessed directly, or through a Network collaboration, iNaturalist data is also distributed to well-known data repositories such as GBIF and ESRI’s Living Atlas and can be easily accessed through their respective portals and data pipelines.

Researchers might assume that regardless of how they access iNaturalist data, their queries return the same sets of observations, the same version of “truth”, but this is not the case. The available data from each platform is nuanced by record obscuration, privacy, and license type, and it is not always immediately clear what data may be unavailable to a researcher through their chosen method of access, nor the unintended downstream effects for researchers who may not be aware of the nuances. This paper explores how using these different datasets in biodiversity research could impact analysis results in unexpected ways. We focused on two main differences in the data contained in the available datasets: how they document record obscuration, and which observation licenses they accept. Using the different versions of iNaturalist observation data, we compare some common types of analysis that are routinely done by researchers and land managers alike, namely environmental niche modeling and species occurrence and spatial range. For our analysis we used both species that iNaturalist obscures under taxon geoprivacy and species it does not. Thus, we assessed whether the pathway through which iNaturalist data are accessed functions as a methodological filter by comparing how platform-specific differences in obscuration metadata and licensing affect spatial coverage, and the bioclimatic variables associated with and represented by occurrence datasets.

## 2. Materials and Methods

### 2.1. Data Sources

We compiled and incorporated details about the data availability from four repositories of iNaturalist data, to quantify and highlight their similarities and differences.

#### 2.1.1. Data directly from iNaturalist

Researchers who want to query iNaturalist data directly, first need a free iNaturalist account. When using the API or the query tools on the website, a researcher can specify exactly the type of data that they need (Callaghan et al. (2026) gives a good overview of using the API). The limitation with this method of data acquisition, apart from the 200,000-record restriction, is that the query does not return the true coordinates, i.e. the location and error as uploaded by the observer, for all records. Instead, where a record has been obscured through either taxon geoprivacy (species has a conservation status), or user geoprivacy (user has restricted the public access), obscured coordinates are returned in place of the true coordinates unless the researcher has specific permission from the observations owner (Rebelo (2020) and iNatHelp (2023b)). These obscured coordinates can be anywhere within roughly a 400 km^2^ grid cell. For more details about record obscuration see iNatHelp (2023a). To highlight the status of a record in their web map, iNaturalist displays obscured records with a semi-transparent circle rather than the opaque locator symbol used for non-obscured records. A researcher can also determine whether a record is obscured or private in the tabular dataset by looking at three fields in the default dataset schema: “geoprivacy” (open| obscured| private), “taxon_geoprivacy” (open| obscured), and “coordinates_obscured” (Boolean). Records that an observer has set as private have no visible coordinates unless the researcher is the owner of them, so will not appear in the webpage map, but can be present in the downloaded data without coordinate or any location information.

#### 2.1.2. Data through a GBIF query

To avoid slowing down their API and to ensure tracking, iNaturalist directs users and researchers to download larger datasets through GBIF (Callaghan et al. (2026)). For researchers, using GBIF is attractive, as the downloads are fast and there are well established computational packages and pipelines for ease of retrieval. Additionally, downloaded GBIF datasets are citable, and contain museum collection records and other credible data sources as well as iNaturalist data. An iNaturalist dataset is provided to GBIF on a weekly basis, therefore the temporal delay of iNaturalist records entering the GBIF dataset is expected to be minimal, compared to getting data from iNaturalist directly, though GBIF may not have the very latest records (iNatHelp (2025b)).

iNaturalist has seven options for licensing records (Anderson (2020)) which can be changed at any time by the account owner. Separate licenses can be applied to observations and media. The current default for all when creating an account is CC BY-NC, but only if users check the box “yes, license my photos, sounds and observations so scientists can use my data”, otherwise it defaults to All rights reserved (iNatHelp (2024)). These available licenses are: Public Domain (CC0), Attribution (CC BY), Attribution NonCommercial (CC BY-NC), Attribution ShareAlike (CC BY-SA), Attribution NonCommercial ShareAlike (CC BY-NC-SA), Attribution NoDerivs (CC BY-ND), Attribution NonCommercial NoDerivs (CC BY-NC-ND), and No License - All Rights Reserved.

GBIF only accepts datasets that are licensed for reuse, so they only accept content with a CC BY or CC BY-NC license, or with the CC0 declaration which releases all property claims over the content so anyone can use it for any reason (gbif.org (2025a) and (gbif.org (2025b)).

GBIF also receives obscured and private records if they meet the Research Grade requirement. If a researcher has downloaded the Darwin Core version of their GBIF query, then the “informationWithheld” field explicitly contains information if the location coordinates were obscured. However, if a researcher downloaded the Simple version of their query, that field is missing ((gbif.org (2025c)). A researcher may infer which records have been obscured from the “coordinateUncertaintyInMeters” field and, to some extent, the “issue” field. However, there is no explicit indicator of obscuration in the simple download and records with a large positional accuracy value may have nothing to do with deliberate obscuration. GBIF datasets contain a direct link to the original iNaturalist record URL which makes it easy to check an individual record at its source.

Another important consideration when integrating biodiversity records from disparate databases is that iNaturalist may not always use the same taxonomy as the GBIF backbone. Cases where a species name is not recognized during the GBIF import may result in that record being rolled up to a higher taxon level (Waller (2022) and Callaghan et al. (2026)) which reduces data resolution and potentially loses the finer level distinction that occurs in the original iNaturalist record. Some examples of taxa missing from the GBIF backbone include *Sciurus brasiliensis* (https://www.inaturalist.org/taxa/1546001-Sciurus-brasiliensis) and *Glocianus punctiger* (https://www.inaturalist.org/taxa/178112-Glocianus-punctiger).

#### 2.1.3. Spatial data through ESRI’s Living Atlas (ArcGIS Online Feature layer)

To access the iNaturalist feature layer in ArcGIS Online fully, a researcher needs an ESRI account. Researchers with access to an ESRI organizational license and who already utilize ESRI products may find this option attractive as for them the data is easily available to pull into their existing mapping workflows and spatial analysis research. The feature layers are available in ESRI’s Living Atlas, which is a curated subset of authoritative data within the broader ArcGIS Online platform. All Living Atlas content is marked as “Authoritative”, meaning it is verified by Esri to be accurate, up-to-date, and ready for use (Guzman (2022)). Therefore, users implicitly trust that the data they see in the iNaturalist feature layer is the same data that they see on iNaturalist.

According to the documentation, ArcGIS Online receives a monthly import of all Research Grade iNaturalist records, except private ones, regardless of the observation license, (ESRI ArcGISOnline (2025), Berry (2024)). Therefore, it is closer in content to an iNaturalist download than obtaining the data through GBIF, though may have a longer lag time.

The iNaturalist feature layer does not distinguish between record coordinates that are true representations and those that have been obscured. Both are symbolized the same way in the displayed data points and there is no clear way within the data attribute table to infer which points are obscured or not. Similar to GBIF’s simple dataset, the positional accuracy field might offer a clue, but a large value may be a result of the observer’s recording device or data entry method, rather than deliberate obscuration. Additionally, none of the original iNaturalist locations fields are in the feature layer attribute table so if a researcher wanted records from a county, for example, they would have to perform a spatial extract using a polygon rather than filter the data. The iNaturalist feature layer does not contain a direct link to the original record in iNaturalist; however, it is possible to link them using the “UUID” field in both the layer and in a separate download from iNaturalist of the same data.

#### 2.1.4. iNaturalist Network Datasets

An iNaturalist Network is a localized iNaturalist website and experience for an individual country, but otherwise connected to the global iNaturalist community. Each Network site is supported by local conservation institutions through formal agreements and are used to promote and benefit local use and biodiversity. These institutions have access to the true coordinates of any observation within their Network (publicly obscured or private) made by an affiliated iNaturalist user. (iNaturalist (2025)).

North America does not have an iNaturalist Network. However, the state of California is trialing a state version of a Network; still with formal agreements in place, but no separate iNaturalist localized website. This “California Data” dataset contains the true coordinates of all taxon geoprivate records, but still the obscured coordinates of user geoprivate records because there is no separate website for users to affiliate with and opt in to. It does not contain private records. Currently this dataset is not available to all researchers, but there are pipelines being built to share the true coordinates with the California Department of Fish and Wildlife through the California Biodiversity Data Exchange (California Academy of Sciences (2026)). The “California Data” was made available to the authors through their work on the project, and was used in some of the analyses for comparison.

### 2.2. Analysis types

#### 2.2.1. Comparing iNaturalist with ARCGIS Online data

To investigate the impact of record obscuration status we compared iNaturalist data with the iNaturalist feature layer in ArcGIS Online (hereafter known as the AGOL dataset, ESRI & iNaturalist (2025)), because of the difficulty identifying obscured records in the latter. As a case study we used observation data for the California Red-legged Frog, *Rana draytonii*. This species was chosen because it is listed as Federally Endangered within North America (U.S. Fish and Wildlife Service (2025)) and is on the IUCN’s Red List (IUCN (2026)) and therefore records are obscured in iNaturalist under taxon geoprivacy. A typical scenario for a land manager with a Threatened and Endangered (T&E) species is to gain understanding of whether the species is present on the landscape and where it has been observed. Species range and coverage also are a consideration for species recovery targets. For instance, in the California Red-legged Frog recovery plan, the criteria for delisting includes knowing that the frogs are successfully re-established in areas of their historic range, that populations of frogs are geographically distributed to allow for migration between individual populations that maintains a viable overall population, and that habitat where populations exist is suitably protected and managed (U.S. Fish and Wildlife Service (2002)).

Data for *R. draytonii* was queried and downloaded from both iNaturalist and AGOL. We investigated the spatial extent and number of observations within counties in the middle of the frog’s current range as an example of a presence analysis within a land management unit. For Figure 1 the iNaturalist data were filtered to obtain records within Marin County using the “county” field, and the AGOL dataset used iNaturalist’s Marin County polygon to perform a spatial clip of records within Marin County. For Table 1, boundary polygons for the nine Bay Area counties were used to spatially clip the data from the AGOL dataset and summarize the number of records per county. The iNaturalist dataset was summarized by the “county” field to get the number of records per county.

**Figure 1:**
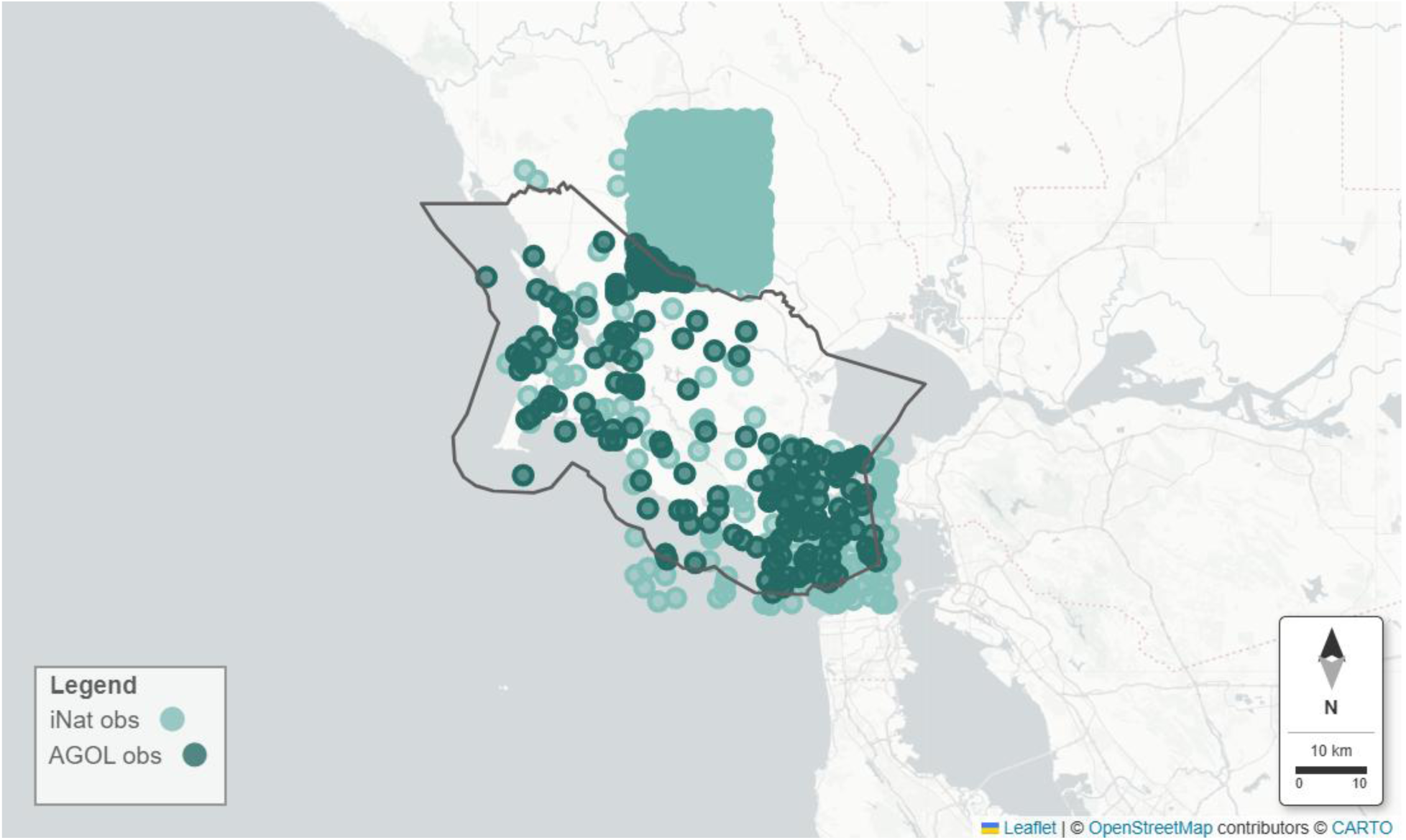
Map of iNaturalist records for *R. draytonii* in Marin County, CA. The light-colored circles are the Marin County observations directly downloaded from iNaturalist, and the darker circles are the same records, but the data is from the AGOL dataset feature layer and “clipped” to the Marin County polygon.

**Table 1:**
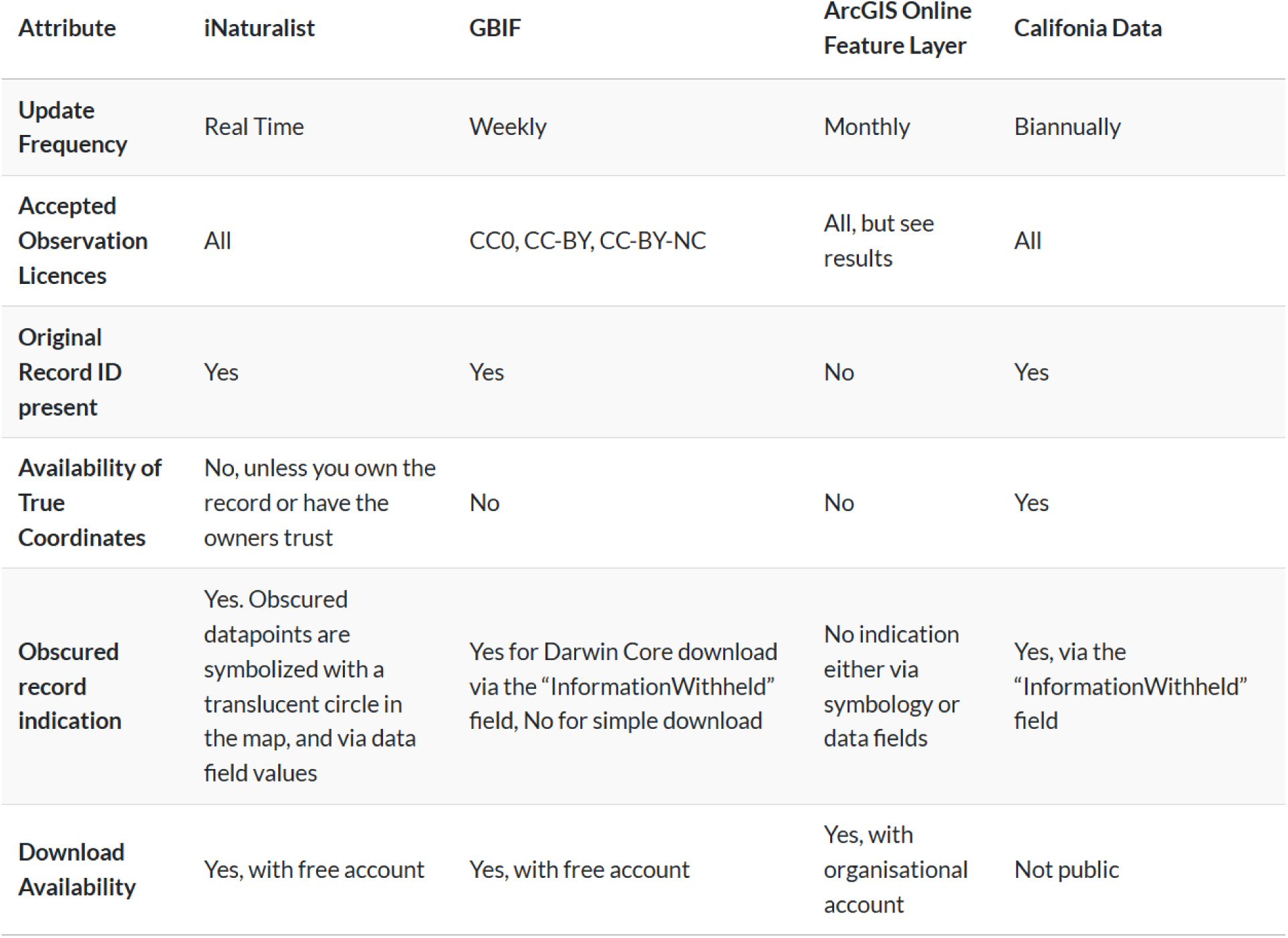
Attribute overview of the different data sources used in this study.

| Attribute | iNaturalist | GBIF | ArcGIS Online<br>Feature Layer | California Data |
| --- | --- | --- | --- | --- |
| Update Frequency | Real Time | Weekly | Monthly | Biannually |
| Accepted Observation Licences | All | CC0, CC-BY, CC-BY-NC | All, but see results | All |
| Original Record ID present | Yes | Yes | No | Yes |
| Availability of True Coordinates | No, unless you own the record or have the owners trust | No | No | Yes |
| Obscured record indication | Yes. Obscured datapoints are symbolized with a translucent circle in the map, and via data field values | Yes for Darwin Core download via the "InformationWithheld" field, No for simple download | No indication either via symbology or data fields | Yes, via the "InformationWithheld" field |
| Download Availability | Yes, with free account | Yes, with free account | Yes, with organisational account | Not public |

#### 2.2.2. Comparing iNaturalist with GBIF data

To investigate the potential downstream effects for ecological analysis of differing license types, we compared the iNaturalist California Data (California Biodiversity Data Exchange et al. (2025)) with GBIF data. As mentioned prior, GBIF only accepts datasets that are licensed with CC0, CC BY or CC BY-NC. For the purpose of this study, we define any license that doesn’t get sent to GBIF as “restrictive”. First, we assessed the proportion of iNaturalist observations associated with restrictive licenses. Second, to understand how “missing” records could have an effect on the spatial distribution of sensitive species with obscured occurrence records, we assessed how the exclusion of restrictively licensed observations for the California Red-legged Frog (*Rana draytonii*) could lead to spatial biases in publicly available occurrence data. Third, we assessed for non sensitive species, how including otherwise ‘missed’ iNaturalist observations from GBIF may alter environmental association comparisons for 19 plant species, particularly with respect to inferred climate associations for ecological forecasting.

Data for the GBIF analyses was prepared in the following ways. To determine the number of restrictive licenses, California Data was filtered to contain Research Grade records only, and summarized. To account for Threatened, Endangered, and rare species (hereafter known collectively as T&E species) in California, we used the California State and Federal lists of Endangered and Threatened Animals and Plants (California Natural Diversity Database (CNDDB). (2025b) and California Natural Diversity Database (CNDDB). (2025a)). The plant list also includes California rare plants. We filtered these lists to include any species with a state code of “SE”, “ST”, or “SR”, or a federal code of “FE” or “FT”. The species list was used to add an identifying column to the data.

To compare spatial extent differences between *R. draytonii* in California, we used a straight GBIF downloaded dataset for *R. draytonii*, (hereafter the GBIF-Direct dataset, GBIF.org User (2025b)) and a modified version of the same GBIF dataset but with all of the iNaturalist data replaced by California Data for *R.draytonii* which contained all iNaturalist Research Grade records regardless of license, and the true coordinates of this sensitive species (hereafter the GBIF-Corrected dataset). We used Uber’s Hexagonal Hierarchical Spatial Index, H3, (Brodsky (2018)) to bin the data points and compare the spatial extent of both datasets. An H3 resolution of 6 was used - each hexagon represents about 36 km^2^ (h3geo.org (2025)).

For environmental niche modeling we used species that are not included in the California State and Federal lists (i.e. non-sensitive species), as we wanted to focus solely on the license issue rather than compounding the results with potential obscuration issues as well. The potential effect of iNaturalist obscured records on niche modeling has been demonstrated already by Contreras-Díaz et al. (2023), where they found that when even a low percentage (25%) of artificially obscured records were present in a dataset of otherwise true coordinates, the species range in both geographic and environmental spaces was significantly expanded, leading to overestimation of ecological niche breadths. The spatial range was restricted to California for analysis efficiency. We used the California Data to select non-sensitive plant species with more than 100 occurrence records. We sorted by the number of restrictive records for each species in descending order, and then selected the top 19 that had a reasonable distribution, i.e. not all 100 observations in the same location. For most of these species, over 50% of records were under restrictive licenses. Similar to the *R draytonii* analysis, two datasets were compiled for each species: a complete dataset of records from GBIF for the 19 species (hereafter, the GBIF-Direct dataset, GBIF.org User (2025a)), and a dataset comprising non-iNaturalist records from GBIF combined with all Research Grade iNaturalist records, regardless of license type, from the California Data (hereafter, the GBIF-Corrected dataset).

In order to ensure that the occurrence data better approximated independent samples of environmental space, we performed spatial thinning to one point per 1km^2^, which reduced observation effort bias and overrepresentation of well surveyed areas. Spatial thinning and niche modelling analyses were conducted in R (R Core Team (2025)) using the packages ecospat (Di Cola et al. (2017) version 4.1.2), terra (Hijmans (2023b) version 1.8-60), and raster (Hijmans (2023a), version 3.6-32).

Environmental data were obtained from the 19 CHELSA v2.1 Bioclimate variables (Karger et al. (2017)), where each distinct climatic parameter related to temperature or precipitation represents annual data averaged over the time period of 1981 to 2010. Bioclimate variables 1–11 correspond to temperature metrics, while variables 12–19 correspond to precipitation metrics. Differences in environmental conditions between the GBIF-Direct and GBIF-Corrected datasets were assessed for each of the 19 Bioclimate variables using the Mann–Whitney U test (Mann & Whitney (1947)), implemented via the wilcox.test() function in R. We used a non-parametric test because the sample sizes were different. Effect sizes were quantified using Cliff’s delta (Cliff (1993)), calculated with the effsize package (Torchiano (2020), version 0.8.1).

## 3. Results

### 3.1. Spatial comparison results with ArcGIS Online Dataset

Observations of *R. draytonii* from Marin County, CA. from iNaturalist and the iNaturalist layer in ArcGIS Online were overlaid to show the effects of spatial clipping on county records. While there are plenty of light-colored iNaturalist data points, and darker colored AGOL dataset points within the county boundary, there is a large square of light colored iNaturalist data points to the north and, to a lesser extent, to the southeast of the county clearly visible in the map. This is an artifact of the obscuration method for geoprivate records, and makes it appear as though the majority of these Marin County records are in neighboring counties. The spatial clipping tool used to query the AGOL dataset records removes any data point outside the polygon boundary, therefore dropping 1362 records from the result. The datapoints within the boundary are only 23.9 percent of the available Marin County records.

Additionally, there are 103 lighter iNaturalist data points that don’t have corresponding darker AGOL points within the Marin County Boundary, which equates to 31.6 percent. A closer examination of the “missing” points within Marin County borders revealed that ArcGIS Online doesn’t seem to receive any records from iNaturalist that have a null value in the license column, indicating that the user chose an “All rights reserved” license for their observations, (77 of the103 missing records). However, there are 26 records that do have licenses, and only 12 of those were created or updated after June 2025, which is when the ArcGIS Online updated the data from iNaturalist used in this dataset. We couldn’t identify the reason why these last 14 points were not included in the AGOL feature layer, which leads to additional concerns in their ingestion method. A researcher may not be aware that they may be missing some Research Grade iNaturalist records when they use data through ArcGIS Online.

Taking the nine California Bay Area counties as a wider sample, there are varying differences in observation numbers between datasets for all except Napa County. Marin and Sonoma Counties have the biggest difference between the number of iNaturalist observations that were actually made in a county (“iNat-County Observations” columns in Table 2 above), and the observation number obtained using a spatial clip of the county boundary (“AGOL-County Clip” column). The differences seem to be most pronounced when there are a lot of observations within a county, or many of the observations are near a boundary. Both of these situations can be observed in Marin County in Figure 1.

**Table 2:**
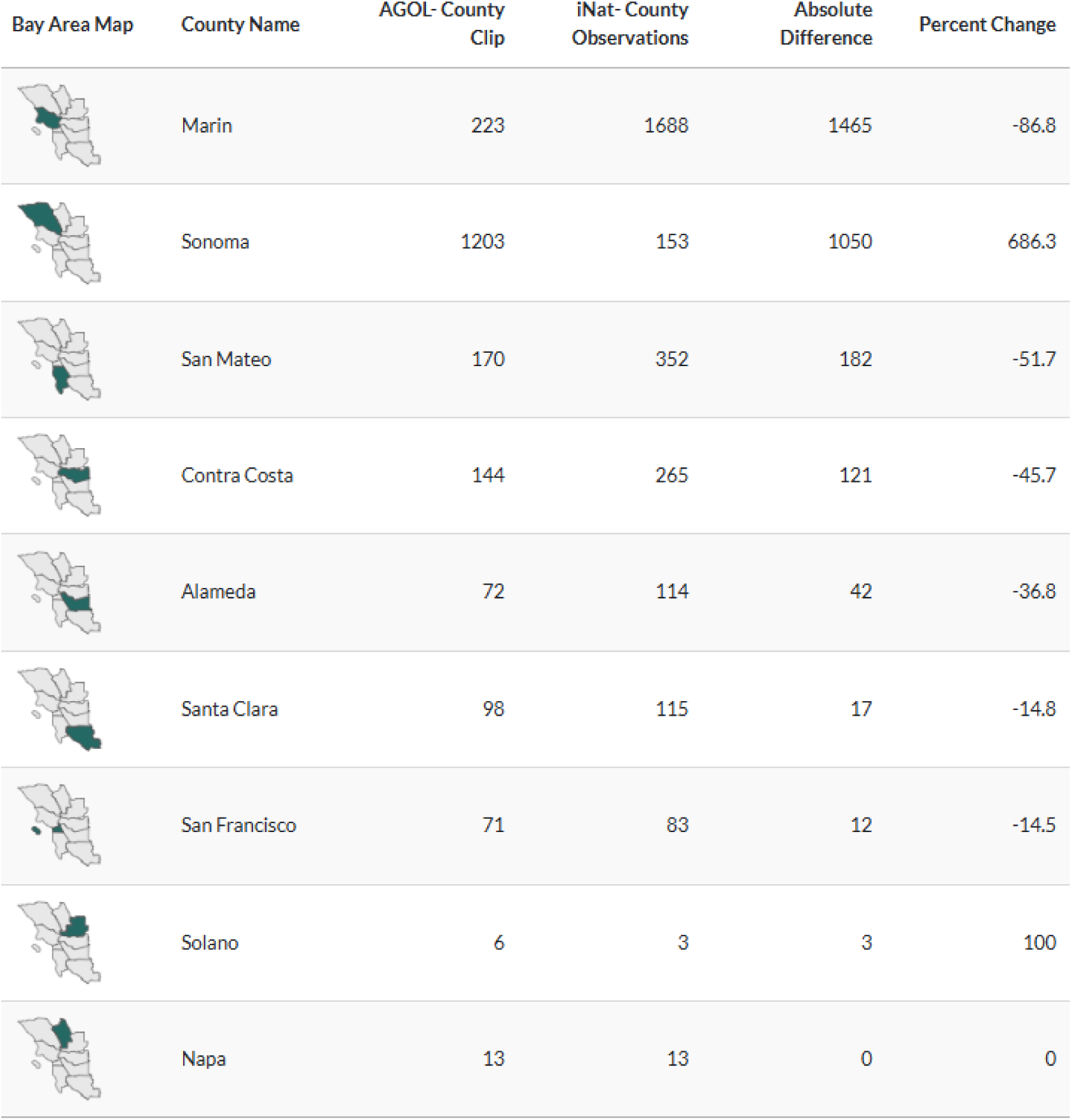
Comparison of iNaturalist filter of the “county” field and performing spatial clips of AGOL feature layer for *R. draytonii* within the nine Bay Area counties in California.

| Bay Area Map | County Name | AGOL- County Clip | iNat- County Observations | Absolute Difference | Percent Change |
| --- | --- | --- | --- | --- | --- |
|  | Marin | 223 | 1688 | 1465 | -86.8 |
|  | Sonoma | 1203 | 153 | 1050 | 686.3 |
|  | San Mateo | 170 | 352 | 182 | -51.7 |
|  | Contra Costa | 144 | 265 | 121 | -45.7 |
|  | Alameda | 72 | 114 | 42 | -36.8 |
|  | Santa Clara | 98 | 115 | 17 | -14.8 |
|  | San Francisco | 71 | 83 | 12 | -14.5 |
|  | Solano | 6 | 3 | 3 | 100 |
|  | Napa | 13 | 13 | 0 | 0 |

### 3.2. License type results with the GBIF dataset

Roughly a quarter (mean = 26.1%) of research grade records in California have restrictive licenses and do not get sent to GBIF. The most striking result in Table 3 (above) is that no clade has fewer than 20% restricted records. That means for each kingdom at least 1 in 5 research grade records do not make it into GBIF purely based on the license type.

**Table 3:**
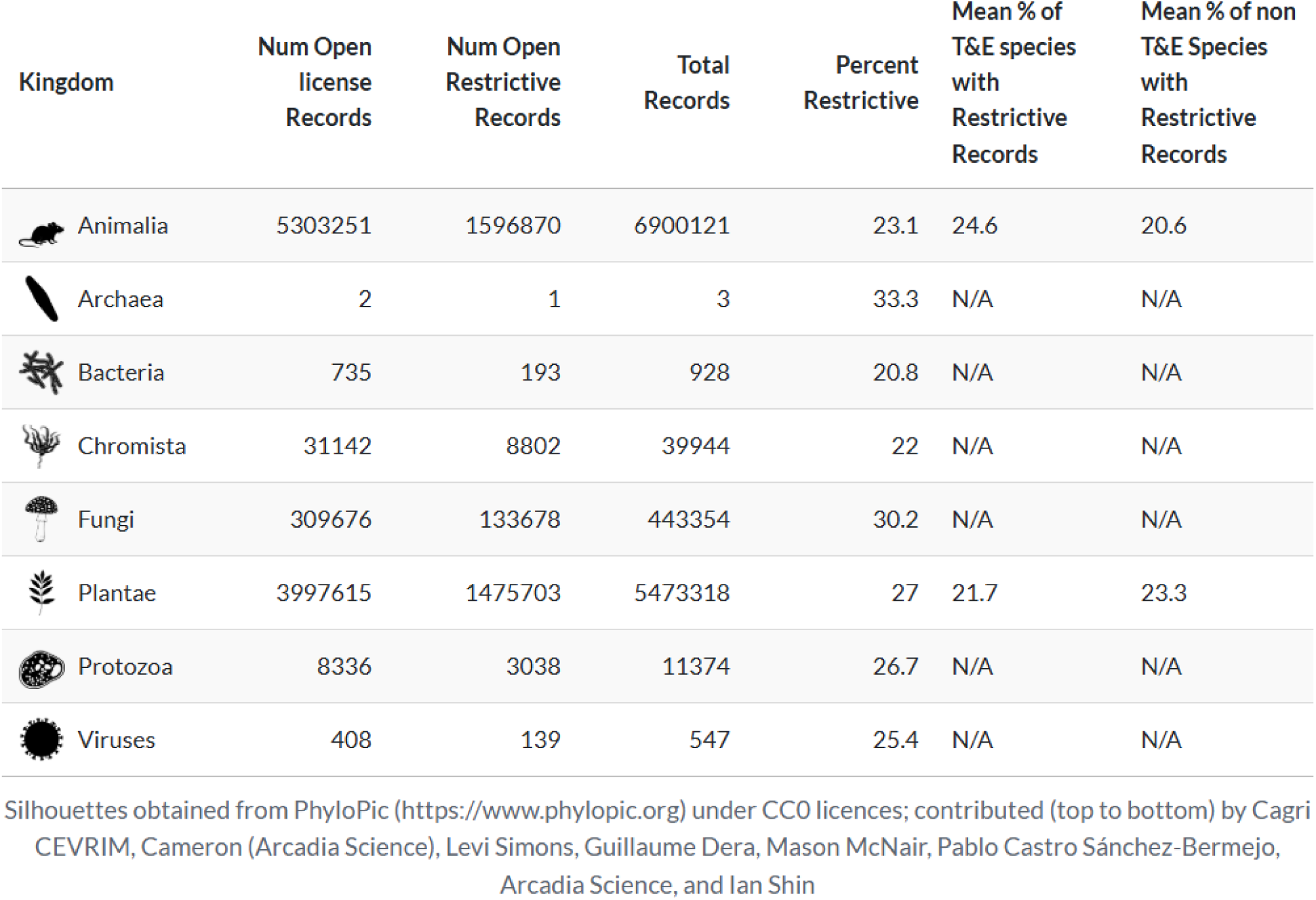
Research grade California iNaturalist records by kingdom, showing the percent of restrictive licenses.

3.2. License type results with the GBIF dataset
| Kingdom | Num Open<br>license<br>Records | Num Open<br>Restrictive<br>Records | Total<br>Records | Percent<br>Restrictive | Mean % of<br>T&E species<br>with<br>Restrictive<br>Records | Mean % of non<br>T&E Species<br>with<br>Restrictive<br>Records |
| --- | --- | --- | --- | --- | --- | --- |
| Animalia | 5303251 | 1596870 | 6900121 | 23.1 | 24.6 | 20.6 |
| Archaea | 2 | 1 | 3 | 33.3 | N/A | N/A |
| Bacteria | 735 | 193 | 928 | 20.8 | N/A | N/A |
| Chromista | 31142 | 8802 | 39944 | 22 | N/A | N/A |
| Fungi | 309676 | 133678 | 443354 | 30.2 | N/A | N/A |
| Plantae | 3997615 | 1475703 | 5473318 | 27 | 21.7 | 23.3 |
| Protozoa | 8336 | 3038 | 11374 | 26.7 | N/A | N/A |
| Viruses | 408 | 139 | 547 | 25.4 | N/A | N/A |

Silhouettes obtained from PhyloPic (https://www.phylopic.org) under CC0 licences; contributed (top to bottom) by Cagri CEVRIM, Cameron (Arcadia Science), Levi Simons, Guillaume Dera, Mason McNair, Pablo Castro Sánchez-Bermejo, Arcadia Science, and Ian Shin

Whilst the percent of restricted records doesn’t differ much between plant and animal T&E species verses non-T&E species, it is important to understand that for some T&E species the percentage is extremely high, with 20 T&E species having 50% or more records with restrictive licenses. This highlights that although the mean percent of restricted records is around a quarter of records, GBIF could be missing a much higher number of iNaturalist records for specific species in its database.

#### 3.2.1. Effects on a sensitive species - Rana draytonii

The combination of obscured records, and restrictive licenses can become compounded and create quite different spatial extents of observations. As mentioned in the methods, data from GBIF (GBIF-Direct dataset) was compared with GBIF data that had all of its iNaturalist records replaced by Research Grade records from the California Data, regardless of license, and with the true coordinates for taxon geoprivate records (GBIF-Corrected dataset).

The false “hot spots” are the dark orange hexagons in Figure 2 (below) which only contain GBIF-Direct observations, and they extend the spatial coverage of *R. draytonii* by 346 cells (out of a total of 853 cells, or 41%). The extension is pronounced around the outside edges of the range particularly to the east and west of the range, but there are some areas where the dark orange hexagons look to be vastly exaggerating the spatial coverage of a local population. This can be seen in the circular inset map where the hexagons with GBIF-Direct observations add a large and false extent to the range to the east of the hexagons containing iNaturalist observations.

**Figure 2:**
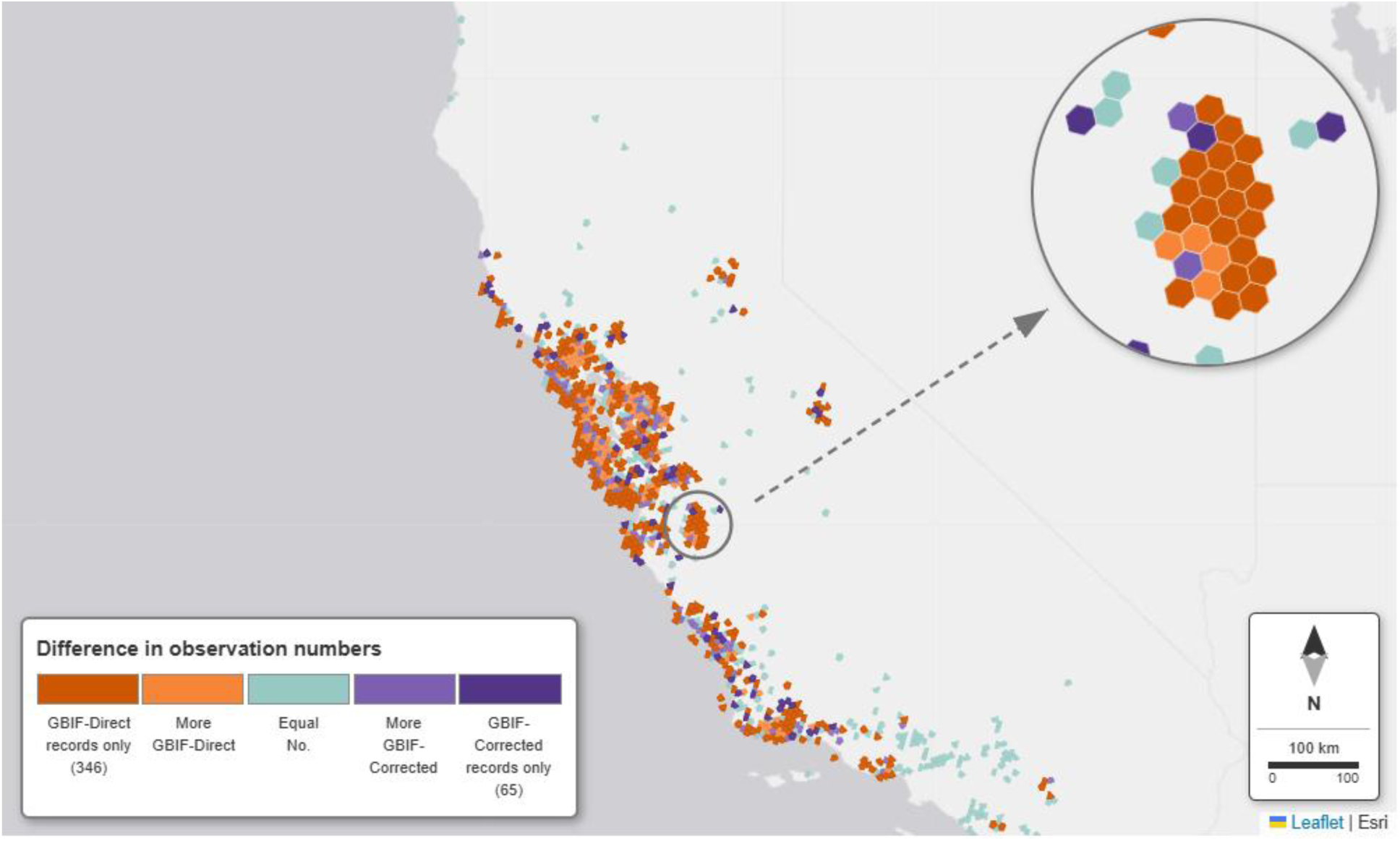
Map of California *R. draytonii* observation numbers within hexagons. The color shows the difference between the number of GBIF-Direct records and GBIF-Corrected in each hexagon; purple contains more GBIF-Corrected records and orange contains more GBIF-Direct. The circular inset map highlights an area with relatively few hexagons containing GBIF-Corrected observations but many with GBIF-Direct only, extending the probable spatial coverage.

By contrast, the false “cold” spots of purple hexagons are areas that only have GBIF-Corrected observations in them. The lack of GBIF-Direct data points in these hexagons make them falsely appear as absent of *R. draytonii* when researchers only use data available through GBIF.

The difference in the number of records between the number of GBIF-Corrected and GBIF-Direct records within hexagons is generally low, with most hexagons having a difference of between −5 and 5, but there was one hexagon that had over 1000 more GBIF-Corrected observations than GBIF-Direct (data point not shown in Figure 3). Out of 442 hexagons with both GBIF-Direct and GBIF-Corrected observations, there were 224 (50.7%) with a difference of zero and it is likely, but not definite, that these hexagons contain the same records from both datasets. Most of the hexagons that contain only GBIF-Direct records (orange bars) have low numbers of observations within them. This makes sense as those records are likely iNaturalist obscured locations in GBIF, and the obscuration process creates spatially randomized points scattered over a large area that may not land within hexagons where frogs were actually observed. Where one hexagon contains many GBIF-Corrected observations, the potential for this wide scattering of obscured records into unoccupied hexagons is much higher. The spatial phenomenon of this can be seen in the circular inset map in Figure 2 where the bottom light purple hexagon actually contains 203 GBIF-Corrected observations. Whilst the random points generated by iNaturalist’s obscuration method likely did place some of those observations within the same hexagon, many appear displaced much further away, giving the larger false range of that population.

**Figure 3:**
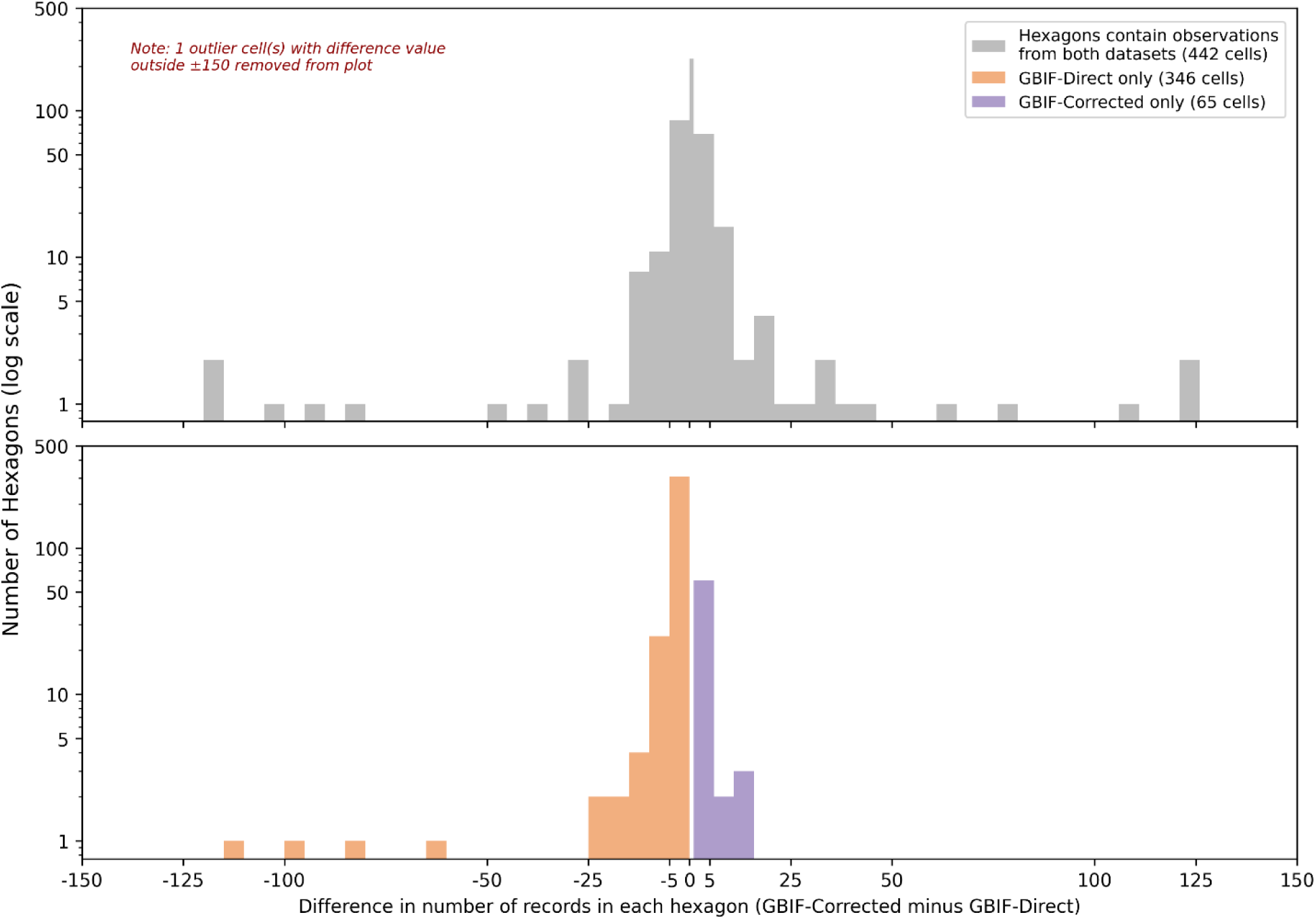
Log scale histogram of difference in the number of records between GBIF-Direct and GBIF-Corrected within hexagon cells. Orange bars represent hexagons with only GBIF-Direct records, and purple bars represent hexagons with only GBIF-Corrected records. Grey bars represent hexagons that contain records from both of the datasets. The overall bin size is 5, except for the zero bin which represents hexagons with equal number of observations from both datasets, i.e. zero difference. The y-axis is on a log scale to better visualize the distribution of difference, especially since many cells have a small difference while a few have large differences, therefore equal visual distances represent order of magnitude differences in the hexagon cell count.

The inclusion of the 720 California Data records for *R. draytonii* that have restrictive licenses (representing 24.0% of *R. draytonii* records in California) as well as the use of true coordinates where they are available, likely contributes to the number of hexagons with more GBIF-Corrected observations than GBIF-Direct, as well as those hexagons with only GBIF-Corrected records. These are the grey bars in Figure 3 that are to the right of the narrow zero bin and the purple bars respectively. Both Figures 2 and 3 demonstrate that at the hexagon scale of 36km^2^, the difference in the number of records between the datasets is not limited to one or two discrepancies, but the majority of cells (74%).

#### 3.2.2. Effects on non-sensitive species

Depending on the species, the lack of iNaturalist restrictive licenses in GBIF has the potential to produce datasets that show a species preference for quite different climatic niche preferences as well as different spatial coverage. In the list of 19 species of plants that we analyzed, most of them had over 50% of their records under restrictive licenses. As stated in the methods, these species were not randomly chosen, they were taken from a list of non-sensitive plant species sorted by their number of restrictive records, but this does highlight that the number of restrictive licenses can vary by species and be much higher than the mean.

When we compared GBIF-Direct and GBIF-Corrected data for environmental niche modeling on these 19 species of plant, only three species showed no real difference in environmental niche using bioclimatic variables (Figure 4 below). Sixteen had at least one bioclimate variable where the difference between the GBIF-Direct and GBIF-Corrected dataset were significant (p < 0.05). Out of those 16 species, four had at least one bioclimate variable where the delta was considered of a medium or large difference. These were *Agave shawii shawii, Argythamnia claryana, Crassula colligata lamprosperma,* and *Lewisia leeana*.

**Figure 4:**
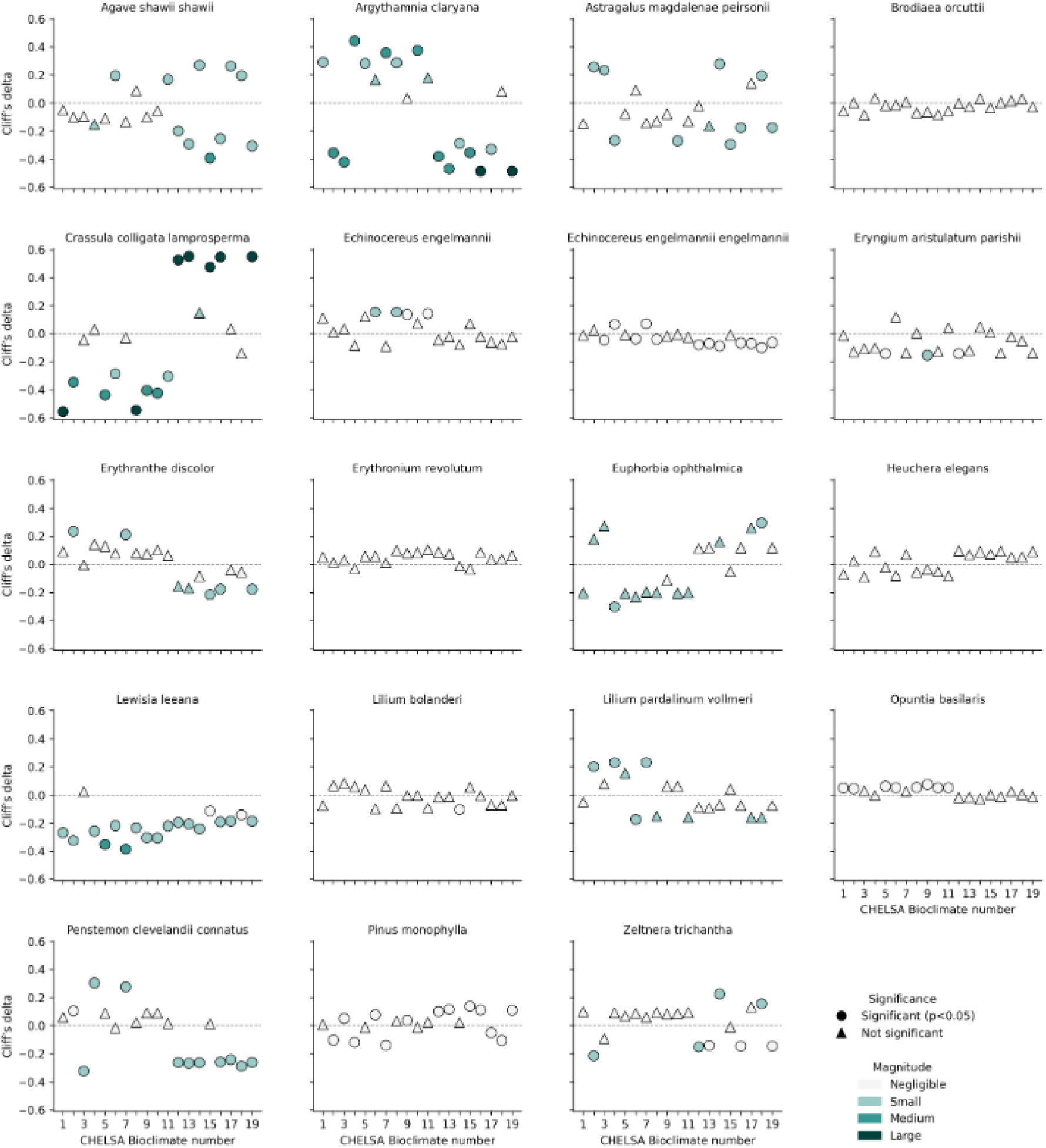
Grid of species charts showing Cliff’s magnitude and delta between the GBIF-Direct and GBIF-Corrected dataset for each of the 19 world bioclimate variables and 19 plant species. The shading represents the magnitude and the symbol represents whether the delta result was significant (p<0.05) indicating that the difference was not due to chance. A positive Cliff’s delta indicates that the GBIF-Corrected values of the bioclimate raster are greater than the GBIF-Direct values. A negative Cliff’s delta indicates that the GBIF-Corrected values of a bioclimate raster are less than the GBIF-Direct values. The larger the magnitude, the more different the two datasets are for that bioclimate variable.

There was a slight indication that temperature bioclimate variables were more affected by the restrictive licenses, but this was very slight (51.1% verses 48.9% of bioclimates with significant deltas), however 2 of the large deltas were for temperature bioclimates and 7 were for precipitation bioclimates (figure 5 above), indicating that GBIF-Corrected modeled niches for some species include populations already thriving in drier conditions, and these were not represented in the GBIF-Direct dataset. The variability of deltas in both temperature and precipitation bioclimates shows that the impact of restrictive licenses on niche modeling is not limited to a specific type of environmental variable, but can affect a range of conditions important for understanding a species’ ecological niche. Models trained on GBIF data alone may overestimate species’ dependence on precipitation, with downstream consequences for climate vulnerability assessments.

**Figure 5:**
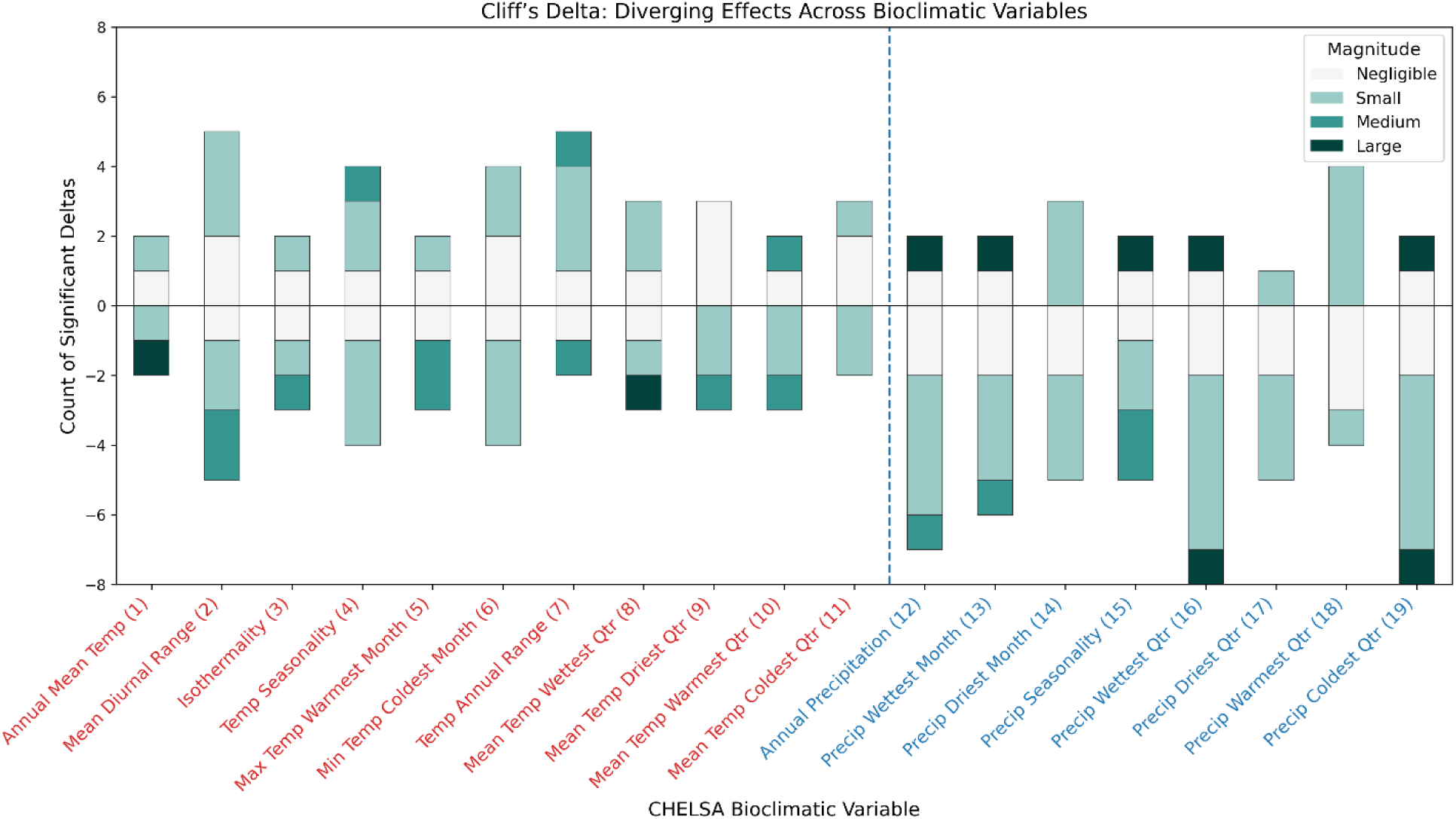
Stacked bar chart showing the count of significant (p<0.05) Cliff’s delta differences between GBIF-Direct and GBIF-Corrected datasets for each of the 19 CHELSA bioclimatic variables across 19 plant species. The bars are colored by the magnitude of the difference, with a dashed line separating temperature-related variables (Bio1–Bio11) from precipitation-related variables (Bio12–Bio19). The x-axis labels are also colored to visually distinguish between temperature and precipitation variables.

## 4. Discussion

Although iNaturalist is a social network for like-minded naturalists (iNaturalist (2026)), it has also become a vast repository of biodiversity data that is being widely used and has unique value for biodiversity research. The influence of iNaturalist data on biodiversity research and monitoring is expected to grow as the platform continues to expand, despite biases in community collected data such as observation effort, taxonomic coverage, spatial, and temporal, that have been discussed at length in previous literature (Courter et al. (2013), Di Cecco et al. (2021), Geurts et al. (2023), Díaz-Calafat et al. (2024), Ellis-Soto et al. (2023), and Callaghan et al. (2026)). Our work in this paper highlights a previously unconsidered source of bias, namely the source from which a researcher obtains their iNaturalist data, and the important implications this can have on analysis. To our knowledge, this is the first study to examine iNaturalist data obtained from these three widely used sources, which are generally assumed to provide the same data, and compare them systematically. Our results highlight that different ways of accessing iNaturalist can function as methodological filters, with measurable consequences for spatial inference, and climatic niche comparisons.

Regardless of whether researchers access data directly from iNaturalist, via ArcGIS Online, or through GBIF, they do not obtain a universal “truth,” (unless they are able to use data directly from a Network) because some records are obscured at the discretion of iNaturalist users and can affect the spatial accuracy of non-sensitive observations. While this limitation does not diminish the importance of iNaturalist as a biodiversity data repository, it underscores the need for researchers to exercise due diligence to ensure that the records they use are fit for their intended purpose. Failure to recognize the differences among these sources, or to select an appropriate version of “truth”, could inadvertently introduce bias into all subsequent analyses.

Our findings highlight important limitations associated with using AGOL iNaturalist data. As seen in the results, researchers interested in geographic boundaries need to spatially clip the AGOL iNaturalist layer, causing discrepancies between the records actually within the area of interest and those whose coordinates are obscured. The impact of these discrepancies could be exacerbated if the county or protected areas of interest are smaller, or there are sensitive species near the boundary. While this issue can affect any species, it is particularly pronounced for sensitive T&E taxa. Some documentation exists regarding obscured records on the iNaturalist feature layer page (ESRI ArcGISOnline (2025)). However, the absence of explicit markers for affected records, whether obscured for taxon geoprivacy or because a user set geoprivacy, makes it impossible for a researcher to assess the extent of the problem within their dataset and fully understand the limitations that this imposes on their analysis.

Additionally, we found that the data iNaturalist provides on a monthly basis to ArcGIS Online do not appear to include all Research Grade observations, as described in the documentation stated. The dataset appears to be missing Research Grade observations that do not have a license; a null in the license field (meaning all rights reserved) appears to prevent the record being sent to the AGOL feature layer, although this exclusion is not clearly documented in the layer metadata. Researchers may be unaware that they are not receiving all of the iNaturalist Research Grade records, and, because it is an authoritative layer in the Living Atlas, may incorrectly assume an exact correspondence with the available iNaturalist data. The absence of direct links to individual iNaturalist records in the AGOL feature layer further complicates efforts to identify missing observations. The number of additional missing Research Grade observations identified in this analysis raises additional questions about why certain records fail to appear in the AGOL layer despite appearing otherwise suitable in iNaturalist.

The limited descriptive metadata associated with these derived datasets can affect not only individual research projects, but also downstream applications that reuse these data for decision support. As a platform, ArcGIS Online is ideally placed for the creation of publicly accessible data aggregator tools, such as dashboards and easy to query applications, based on spatial boundaries such as counties or land parcels. Because these tools are often designed for users without specialized subject matter expertise on biodiversity informatics, insufficient metadata can lead to overconfident interpretation of spatial outputs. Although ArcGIS Online makes it straightforward to discover and incorporate spatial layers into dashboards, applications, and other decision-support tools, the available metadata may be insufficient for developers to determine whether a layer is appropriate for the intended analytical use.

We therefore recommend iNaturalist and ArcGIS Online introduce a higher degree of clarity in the AGOL feature layers descriptive metadata, stating exactly which records are included, as well as providing explicit markers for obscured records in the form of both symbology and attribute table fields. Improving transparency between iNaturalist data and the AGOL feature layer could be achieved by including the record level iNaturalist URLs in the AGOL attribute table. Because iNaturalist already exports data to GBIF using the Darwin Core data schema, which contains these fields as well as other data quality fields, incorporating all comparable fields into the AGOL feature layer would improve transparency and record verification for an ArcGIS Online export. This could reduce the number of potential issues with using AGOL iNaturalist data.

iNaturalist has a dedicated community of voluntary identifiers who use their collective expertise to turn observations from casual to Research Grade, and making those records the most useful to science. 57.2% of all iNaturalist records make it to Research Grade within one year of their recorded date, and the rate of conversion has been increasing each year (Campbell et al. (2023)). Not all observations can achieve Research Grade however, they may be of a cultivated organism, or of poor identification quality. At the time of writing, iNaturalist currently has 311,686,520 observations, with 175,190,115 having obtained Research Grade (56.2%) which is not far off the percentage that Cambel et all found, indicating that just over half of iNaturalist observations have the ability to ever reach Research Grade and that the identifiers are keeping up with the influx of observations. Our GBIF comparison analysis into restrictive licenses showed that for California at least, 26.1% of CA Research Grade observations will not be exported to GBIF due to having restrictive licenses. If similar licensing patterns apply globally, we see that potentially GBIF could be missing out on 45,668,537 records. Whilst it is acknowledged that not all of these “missing” Research Grade observations will be valuable to GBIF users, this can constitute a lot of information to not be available, without full assessments of how the missing data may bias an analysis.

Our analysis highlighted a compounding effect between taxon obscured records entering GBIF, and restrictive license records not entering GBIF. The hot and cold spots in the *Rana draytonii* map provides a much larger spatial extent when using GBIF data, than with the full complement of iNaturalist records. They also have some missing values where records actually exist. As a large proportion of these “false” cells are around the edges of the spatial extent, this could cause interpretation issues with research into species State recovery targets, as it may provide false evidence of species range expansion or falsely pass a recovery threshold.

We also showed that for any species, niche modeling results may be different when comparing the data downloaded from GBIF with what the dataset could look like if iNaturalist records were not restricted. This effect would depend upon the proportion of iNaturalist records for a specific species in GBIF and then the proportion of those records that are not included due to restrictive licenses. However, it is an effect that researchers should be aware of when using GBIF data for environmental niche modeling. Our analysis used a course resolution predictor as an example, but finer scale predictors from SDMs which allow for study of microclimates for example could show even greater disparities. iNaturalist is a major contributor to GBIF (Mason et al. (2025)), and while the total amount of iNaturalist records in GBIF is less than 5% overall (Xue (2025)), for recent records it is more influential, with 42% of GBIF species having 100% of their records provided by iNaturalist between 2020 and 2023 (Loarie (2023)). The proportion of modern records from iNaturalist could become particularly problematic for research in specialized species (where there are very few records), or areas with restricted access, where iNaturalist observers may feel more inclined to use greater license restrictions, such as on their own property.

Across many of the 19 plant species we analyzed, we found that the “missing” iNaturalist records made the environmental niches created with the GBIF dataset appear to be areas of higher annual precipitation (Bio12) and wetter-quarter precipitation (Bio16) than with the additions. This is potentially ecologically meaningful as it could, for example, lead to mis-identifying which species populations are most vulnerable to climate change, or which restoration seed zones are appropriate for collection. However, it also suggests a potential observer bias of who is applying restrictive licenses to their records.

We can’t untangle the effects of observer bias when interpreting our results, but it is an interesting avenue for future research to understand who is applying restrictive licenses and why, and how this may affect the data that is available for research. One possible explanation is that these observers may be more experienced botanists, professional ecologists, and dedicated naturalists who are often more intentional about their data. They’re more likely to be doing targeted botanical surveys rather than casual opportunistic recording. In California, it may be that such surveys disproportionately fall within harsher climates such as dry interior foothill and montane habitats or remote desert-adjacent zones, and many other areas away from established urban places that casual observers simply don’t reach.

We recommend a two-pronged approach for improving the data quality and comparability in the flow between iNaturalist and GBIF. The first approach is to address the record licensing. While we acknowledge that all iNaturalist users should retain the ability to choose the level of licensing that they are comfortable with, we suggest it needs to be made extremely clear in iNaturalist what the user is giving up by not making their data as useful to research when selecting a restrictive observation license, and show how the terms of the more restrictive licenses apply to the data. The language provided when a user creates an account is a start, but it could be much more robust and explicit. For instance, deselecting the box for “please license my…”, sets the default license as all rights reserved (the most restrictive license), without going into details about all the options. Unlike other datasets submitted to GBIF, iNaturalist data is not static, and any change made to an iNaturalist record, such as the identification or license type gets reflected in the next data submission to GBIF. This can make data disappear if, for instance, an iNaturalist user deletes their account, but it also means that if current users with restrictive licenses can be encouraged to change to a more open license, all of their Research Grade records become available to GBIF with very little effort. When surveyed, participatory scientists rate making meaningful contributions to science and conservation as one of their main reasons for participating. It ranked important or very important for 99% of FrogID citizen science project survey respondents (Thompson et al. (2023)), and was the primary motivating reason for 40% of Christmas Bird Count survey respondents, and the continuing motivation for 55% (Larson et al. (2020)). So it may be that some iNaturalist users are unaware that their choice of license for observations and media affect downstream data reuse for biodiversity science. Organizers of community science events could include language around licensing in their promotional outreach to remind attendees that their observations are being widely used by the research community and their observation licensing choice really matters downstream.

The second suggestion is to address the obscured records in GBIF itself, by making them explicit in every GBIF data download. Making sure all obtainable versions of GBIF data have the same important information, including the ‘informationWithheld’ field, would allow all researchers to make informed decisions about whether the data resolution is suitable for their analysis or not.

Our comparison revealed substantial differences in the datasets from iNaturalist, AGOL, and GBIF, and potential effects on even simple data analysis. Although these differences are lightly mentioned in each dataset’s metadata, the ramifications of using one dataset over another for analysis are not. This paper does not seek to diminish the considerable benefits of utilizing iNaturalist and other participatory science in biodiversity analysis, rather it intends to daylight another consideration researchers need to make before conducting any form of analysis. We therefore suggest that researchers thoroughly consider and investigate each dataset carefully and make sure it fits their research needs. Because AGOL does not clearly identify obscured iNaturalist coordinates, we recommend that iNaturalist data accessed through AGOL be used only for coarser-scale spatial analyses and not for determining whether a species is absent from a given parcel of land. When researchers use GBIF data, we suggest that they supplement, if feasible, the “missing” data from iNaturalist directly to ensure that they have the most complete data available to them, providing that the research fits within the limitations of all of the Creative Commons licenses. Finally, for complete scientific transparency and reproducibility, any analysis, paper, or application using iNaturalist data should state exactly which dataset was used and consider potential additional biases this may have introduced. Our results also reinforce a broader point: biodiversity datasets are shaped not only by ecological processes and platform-specific rules, but also by underlying social (and economic structures) such as an individual’s right to choose what license to give their observation, and these forces can propagate into conservation assessments (Ellis-Soto et al. (2024)).

## Notes

### Competing Interest Statement

The authors have declared no competing interest.

## References

1. Anderson, E. (2020). Tech Tip Tuesday: Understanding Licensing. In iNaturalist. https://www.inaturalist.org/posts/30692-tech-tip-tuesday-understanding-licensing; iNaturalist.

2. Backstrom, L. J., Callaghan, C. T., Leseberg, N. P., Sanderson, C., Fuller, R. A., & Watson, J. E. M. (2024). Assessing adequacy of citizen science datasets for biodiversity monitoring. Ecology and Evolution, 14(2), e10857. 10.1002/ece3.10857

3. Berry, L. (2024). iNaturalist Observations Now Available within ArcGIS Living Atlas (Beta Release). In ArcGIS Blog. https://www.esri.com/arcgis-blog/products/arcgis-living-atlas/announcements/inaturalist-living-atlas-beta-release.

4. Brodsky, I. (2018). H3: Uber’s Hexagonal Hierarchical Spatial Index. In Uber Blog. https://www.uber.com/en-EG/blog/h3/.

5. California Academy of Sciences. (2026). California Biodiversity Data Exchange - California Academy of Sciences. In California Biodiversity Data Exchange. https://www.calacademy.org/community-science/california-biodiversity-data-exchange.

6. California Biodiversity Data Exchange, iNaturalist, & California Academy of Sciences. (2025). The California Data.

7. California Natural Diversity Database (CNDDB). (2025a). State and Federally Listed Endangered and Threatened Animals of California. California Department of Fish and Wildlife. Sacramento, CA.

8. California Natural Diversity Database (CNDDB). (2025b). State and Federally Listed Endangered, Threatened, and Rare Plants of California. California Department of Fish and Wildlife. Sacramento, CA.

9. Callaghan, C. T., Campbell, C. J., Widness, J., Goldstein, B. R., Mesaglio, T., Seltzer, C., Iwane, T., Guralnick, R., & Mason, B. M. (2026). Guidelines and best practices for the scientific use of global iNaturalist data. EcoEvoRxiv.

10. Callaghan, C. T., Winnebald, C., Smith, B., Mason, B. M., & López-Hoffman, L. (2025). Citizen science as a valuable tool for environmental review. Frontiers in Ecology and the Environment, 23(1), e2808. 10.1002/fee.2808

11. Campbell, C. J., Barve, V., Belitz, M. W., Doby, J. R., White, E., Seltzer, C., Di Cecco, G., Hurlbert, A. H., & Guralnick, R. (2023). Identifying the identifiers: How iNaturalist facilitates collaborative, research-relevant data generation and why it matters for biodiversity science. BioScience, 73(7), 533–541. 10.1093/biosci/biad051

12. CBD. (2022). Report of the conference of the parties to the convention on biological diversity on the second part of its fifteenth meeting (Kunming-{{Montreal Global Biodiversity Framework}}, {{Draft Decision Submitted}} by the {{President}}. CBD/COP/15/L.25 2022).

13. Cliff, N. (1993). Dominance statistics: Ordinal analyses to answer ordinal questions. Psychological Bulletin, 114(3), 494–509.

14. Contreras-Díaz, R. G., Nori, J., Chiappa-Carrara, X., Peterson, A. T., Soberón, J., & Osorio-Olvera, L. (2023). Well-intentioned initiatives hinder understanding biodiversity conservation: Cloaked iNaturalist information for threatened species. Biological Conservation, 282, 110042. 10.1016/j.biocon.2023.110042

15. Courter, J. R., Johnson, R. J., Stuyck, C. M., Lang, B. A., & Kaiser, E. W. (2013). Weekend bias in Citizen Science data reporting: Implications for phenology studies. International Journal of Biometeorology, 57(5), 715–720. 10.1007/s00484-012-0598-7

16. Di Cecco, G. J., Barve, V., Belitz, M. W., Stucky, B. J., Guralnick, R. P., & Hurlbert, A. H. (2021). Observing the Observers: How Participants Contribute Data to iNaturalist and Implications for Biodiversity Science. BioScience, 71(11), 1179–1188. 10.1093/biosci/biab093

17. Di Cola, V., Broennimann, O., Petitpierre, B., Breiner, F. T., D’Amen, M., Randin, C., Engler, R., Pottier, J., Pio, D., Dubuis, A., Pellissier, L., Mateo, R. G., Hordijk, W., Salamin, N., & Guisan, A. (2017). Ecospat: An R package to support spatial analyses and modeling of species niches and distributions. Ecography, 40(6), 774–787. 10.1111/ecog.02671

18. Díaz-Calafat, J., Jaume-Ramis, S., Soacha, K., Álvarez, A., & Piera, J. (2024). Revealing biases in insect observations: A comparative analysis between academic and citizen science data. PLOS ONE, 19(7), e0305757. 10.1371/journal.pone.0305757

19. Ellis-Soto, D., Chapman, M., & Koltz, A. M. (2024). Addressing data disparities is critical for biodiversity assessments. Trends in Ecology & Evolution, 39(12), 1066–1069. 10.1016/j.tree.2024.10.005

20. Ellis-Soto, D., Chapman, M., & Locke, D. H. (2023). Historical redlining is associated with increasing geographical disparities in bird biodiversity sampling in the United States. Nature Human Behaviour, 7(11), 1869–1877. 10.1038/s41562-023-01688-5

21. ESRI ArcGIS Online. (2025). iNaturalist Observations - Overview.

22. ESRI, & iNaturalist. (2025). iNaturalist Observations [Feature Layer].

23. gbif.org. (2025a). Quick guide to publishing data through GBIF.org. In gbif.org. https://www.gbif.org/publishing-data.

24. gbif.org. (2025b). iNaturalist Research-grade Observations. https://www.gbif.org/dataset/50c9509d-22c7-4a22-a47d-8c48425ef4a7.

25. gbif.org. (2025c). Occurance download formats. In Technical Documentation. https://techdocs.gbif.org/en/data-use/download-formats.

26. GBIF.org User. (2025a). Plant Species DarwinCore download. The Global Biodiversity Information Facility. 10.15468/DL.PKPQAF

27. GBIF.org User. (2025b). R.Draytonii Simple download. The Global Biodiversity Information Facility. 10.15468/DL.7ABD3G

28. Geurts, E. M., Reynolds, J. D., & Starzomski, B. M. (2023). Turning observations into biodiversity data: Broadscale spatial biases in community science. Ecosphere, 14(6), e4582. 10.1002/ecs2.4582

29. Güntsch, A., Overmann, J., Ebert, B., Bonn, A., Le Bras, Y., Engel, T., Hovstad, K. A., Lange Canhos, D. A., Newman, P., van Ommen Kloeke, E., Ratcliffe, S., le Roux, M., Smith, V. S., Triebel, D., Fichtmueller, D., & Luther, K. (2024). National biodiversity data infrastructures: Ten essential functions for science, policy, and practice. 75(2), 139–151.

30. Guzman, L. D. (2022). Living Atlas - Customer FAQ’s. In ArcGIS StoryMaps. https://storymaps.arcgis.com/stories/e40b456d926941b3ba18b383fe0ec8c8.

31. h3geo.org. (2025). Tables of Cell Statistics Across Resolutions H3. In H3 docs. https://h3geo.org/docs/core-library/restable/.

32. Hijmans, R. J. (2023a). Raster: Geographic data analysis and modeling [Manual].

33. Hijmans, R. J. (2023b). Terra: Spatial data analysis [Manual].

34. iNatHelp. (2023a). What is geoprivacy? What does it mean for an observation to be obscured? In iNaturalist Help. https://help.inaturalist.org/en/support/solutions/articles/151000169938-what-is-geoprivacy-what-does-it-mean-for-an-observation-to-be-obscured-.

35. iNatHelp. (2023b). How can I access the hidden coordinates of sensitive species? In iNaturalist Help. https://help.inaturalist.org/en/support/solutions/articles/151000170343-how-can-i-access-the-hidden-coordinates-of-sensitive-species-.

36. iNatHelp. (2024). How to Sign Up for an iNaturalist Account. In iNaturalist Help. https://help.inaturalist.org/en/support/solutions/articles/151000195690-how-to-sign-up-for-an-inaturalist-account.

37. iNatHelp. (2025a). What is the Data Quality Assessment and how do observations qualify to become “Research Grade”. In iNaturalist Help. https://help.inaturalist.org/en/support/solutions/articles/151000169936-what-is-the-data-quality-assessment-and-how-do-observations-qualify-to-become-research-grade-.

38. iNatHelp. (2025b). Which iNaturalist observations are exported for GBIF, and how often does this export happen? In iNaturalist Help. https://help.inaturalist.org/en/support/solutions/articles/151000170346-which-inaturalist-observations-are-exported-for-gbif-and-how-often-does-this-export-happen-.

39. iNaturalist. (2025). iNaturalist Network. In iNaturalist. https://www.inaturalist.org/sites/network.

40. iNaturalist. (2026). About iNaturalist. In iNaturalist. https://www.inaturalist.org/pages/about.

41. IPBES. (2019). Global assessment report on biodiversity and ecosystem services of the Intergovernmental Science-Policy Platform on Biodiversity and Ecosystem Services. Zenodo. 10.5281/zenodo.6417333

42. IUCN. (2026). California Red-legged Frog. In IUCN Red List of Threatened Species. https://www.iucnredlist.org/en.

43. Karger, D. N., Conrad, O., Böhner, J., Kawohl, T., Kreft, H., Soria-Auza, R. W., Zimmermann, N. E., Linder, H. P., & Kessler, M. (2017). Climatologies at high resolution for the earth’s land surface areas. Scientific Data, 4(1), 170122. 10.1038/sdata.2017.122

44. Larson, L. R., Cooper, C. B., Futch, S., Singh, D., Shipley, N. J., Dale, K., LeBaron, G. S., & Takekawa, J. Y. (2020). The diverse motivations of citizen scientists: Does conservation emphasis grow as volunteer participation progresses? Biological Conservation, 242, 108428. 10.1016/j.biocon.2020.108428

45. Loarie, S. (2023). Thank you for helping generate most GBIF records for most species since 2020. In iNaturalist. https://www.inaturalist.org/blog/76606-thank-you-for-helping-generate-most-gbif-records-for-most-species-since-2020; iNaturalist.

46. Mann, H. B., & Whitney, D. R. (1947). On a test of whether one of two random variables is stochastically larger than the other. Annals of Mathematical Statistics, 18(1), 50–60.

47. Mason, B. M., Mesaglio, T., Barratt Heitmann, J., Chandler, M., Chowdhury, S., Gorta, S. B. Z., Grattarola, F., Groom, Q., Hitchcock, C., Hoskins, L., Lowe, S. K., Marquis, M., Pernat, N., Shirey, V., Baasanmunkh, S., & Callaghan, C. T. (2025). iNaturalist accelerates biodiversity research. BioScience, biaf104. 10.1093/biosci/biaf104

48. R Core Team. (2025). R: A language and environment for statistical computing [Manual]. R Foundation for Statistical Computing.

49. Rebelo, T. (2020). Using iNaturalist data for research. In iNaturalist. https://www.inaturalist.org/posts/44352-using-inaturalist-data-for-research; iNaturalist.

50. Secretariat of the Convention on Biological Diversity. (2024a). 2030 Targets (with Guidance Notes). In Kunming-Montreal Globval Biodiversity Framework. https://www.cbd.int/gbf/targets; Secretariat of the Convention on Biological Diversity.

51. Secretariat of the Convention on Biological Diversity. (2024b). Kunming-Montreal Global Biodiversity Framework. In Kunming-Montreal Globval Biodiversity Framework. https://www.cbd.int/gbf; Secretariat of the Convention on Biological Diversity.

52. Thompson, M. M., Moon, K., Woods, A., Rowley, J. J. L., Poore, A. G. B., Kingsford, R. T., & Callaghan, C. T. (2023). Citizen science participant motivations and behaviour: Implications for biodiversity data coverage. Biological Conservation, 282, 110079. 10.1016/j.biocon.2023.110079

53. Torchiano, M. (2020). Effsize: Efficient effect size computation [Manual].

54. U.S. Fish and Wildlife Service. (2002). Recovery Plan for the California Red-legged Frog (Rana aurora draytonii) (p. 180). U.S. Fish and Wildlife Service.

55. U.S. Fish and Wildlife Service. (2025). California Red-legged Frog (Rana draytonii). https://www.fws.gov/species/california-red-legged-frog-rana-draytonii.

56. Waller, J. T. (2022). Finding Data Gaps in the GBIF Backbone Taxonomy. Biodiversity Information Science and Standards, 6, e91312. 10.3897/biss.6.91312

57. Xue, L. (2025). Understanding how iNaturalist data affect GBIF and its limitations: A guide to key considerations. https://open.library.ubc.ca/soa/cIRcle/collections/undergraduateresearch/52966/items/1.0449748. 10.14288/1.0449748

